# Species delimitation analysis indicates cryptic speciation for *Terpios gelatinosus* (Porifera, Demospongiae) from coastal regions of the northeastern Mediterranean Sea

**DOI:** 10.1101/2024.01.12.575186

**Authors:** Esra Öztürk, Berivan Temiz, Arzu Karahan

## Abstract

Sponges, comprising over 9000 recorded species, represent one of the most primitive groups of metazoans. Given the importance of species distribution records and the identification of new species in scientific research, these endeavors play a crucial role in enhancing ecological insights, conserving biodiversity, facilitating a better understanding of the relationships between various groups of organisms, advancing our knowledge of evolution, and potentially expanding biomedical implications. In this study, we used basic morphological data, mitochondrial Cytochrome Oxidase I (COI) gene, and Internal Transcribed Spacer two (ITS2) regions to evaluate the *Terpios* samples collected from four coastal sites along the northeastern Mediterranean Sea, covering a distance of approximately 450 km. For this purpose, 81 COI and 11 ITS2 records of the order Suberitida were mined from GenBank, and species delimitation analysis was performed using both the Automatic Barcode Gap Discovery method and the Poisson Tree Process (PTP), along with 11 samples from the present study. While we observed slight differences in spicule sizes, the overall morphology of all samples from our study was similar. Within the scope of this research, we report the first-ever presence record of *T. gelatinosus* in the northeastern Mediterranean Sea. Furthermore, we document evidence pointing to the potential existence of cryptic speciation in the region.

## 1. Introduction

Among metazoans, sponges (Porifera) are the most primitive group, with 9,697 species known to date (Müller *et al*., 2007; de Voogd *et al*., 2024). Calcarea, Demospongiae, Hexactinellida, and Homoscleromorpha are the four classes into which they are divided (van Soest *et al*., 2012). With about 8,000 species worldwide, the largest class of Porifera is Demospongiae, which includes almost 90% of all living sponges (Lavrov *et al*., 2023). From the tropics to the polar seas, they inhabit a variety of habitats and substrates (Wörheide *et al*., 2012). The eastern Mediterranean Sea has a high species abundance because of its diverse and suitable rocky shorelines (Gerovasileiou & Voultsiadou-Koukoura, 2012).

For about three centuries, species classification has been done using morphology-based taxonomy (Dunn, 2003). However, the identification of primitive marine animals based on only classical taxonomy has been an ongoing problem due to cryptic speciation, convergent evolution, and simple and non-uniform body shape (Solé-Cava & Thorpe, 1986; Park *et al*., 2007; Morrow *et al*., 2013). One of the main morphological characteristics of sponges is spicules, but most of them are quite simple, and their size can easily change even under relatively low environmental variation (Solé-Cava & Thorpe, 1986). Thus, molecular markers were suggested to support morphology-based taxonomy (Solé-Cava *et al*., 1991). DNA barcoding has been used for this purpose for around two decades (Hebert *et al*., 2003a; Hebert & Gregory, 2005). Over a million species have been barcoded by now (Ratnasingham and Hebert, 2007), and this number is increasing daily. Besides, there are certain DNA barcoding initiatives that focus on only specific marine groups, like the ‘Sponge Barcoding Project’ (Vargas *et al*., 2012; Wörheide and Erpenbeck, 2007). The Cytochrome Oxidase I gene is the primary gene region used for the barcoding purpose (Hebert *et al*., 2003b; Karahan *et al*., 2017); the Internal Transcribed Spacer (ITS) region (Wörheide *et al*., 2004) and many others (van Oppen *et al*., 2002) are used for different taxa to delimitate the species.

The genus *Terpios*, classified under the Suberitidae family of Demospongiae, is found under diverse temperatures and habitats, from animal shells to the coastal and deep waters (van Soest, 2002, *et al*., 2012). *Terpios gelatinosus* (Bowerbank, 1866) displays a light brownish yellow or bright brown color. It has been reported that the color alters to deep blue in the presence of a symbiotic alga (Ackers *et al*., 2007). So far, *T. gelatinosus* records have been given in Italy (Bertolino *et al*., 2013), Portugal (Monteiro, 2013), Greece (Voultsiadou-Koukoura *et al*., 2016), and the Mediterranean coast of Israel, with a single public DNA barcode record in the database (Idan *et al*., 2018). The species was also reported in the eastern Aegean Sea (Topaloğlu & Evcen, 2014).

This study aimed to map the distribution of *Terpios gelatinosus* along the northeastern coasts of Türkiye, covering a distance of 450 km. This was accomplished via species delimitation analysis (SDA), utilizing two molecular markers (COI and ITS2), alongside an assessment of basic morphological traits. Furthermore, we incorporated the database-mined samples into our analysis to augment our comprehension of the species.

## 2. Materials and Methods

Eleven *Terpios* samples were obtained from rocky coastal regions with a depth of under 1 meter between November 2017 and October 2018, spanning 450 km of the Turkish Mediterranean coastline, encompassing three locations in Mersin (Mezitli (n=2), Kızkalesi (n=2), and Tisan (n=4)) and one location in Antalya (Side (n=3)) (Figure 1). Photographs of all specimens were captured in their natural habitats (beneath stones), and their colors and overall morphologies were documented. A fragment from each specimen was removed from the surface using a razor blade and placed into a 1.5-ml vial for DNA extraction (Paz et al., 2003). And also, a fragment of an individual from each site was preserved in formalin (10%) for morphological examination. Spicules were prepared for microscopy following the protocol of Schwab and Shore (1971), and basic morphological analysis was conducted using stereo and light microscopes (Olympus SZX16-UC30 and Olympus CX43-ToupTek cameras) in accordance with Bowerbank’s criteria (1866).

**Figure 1.**
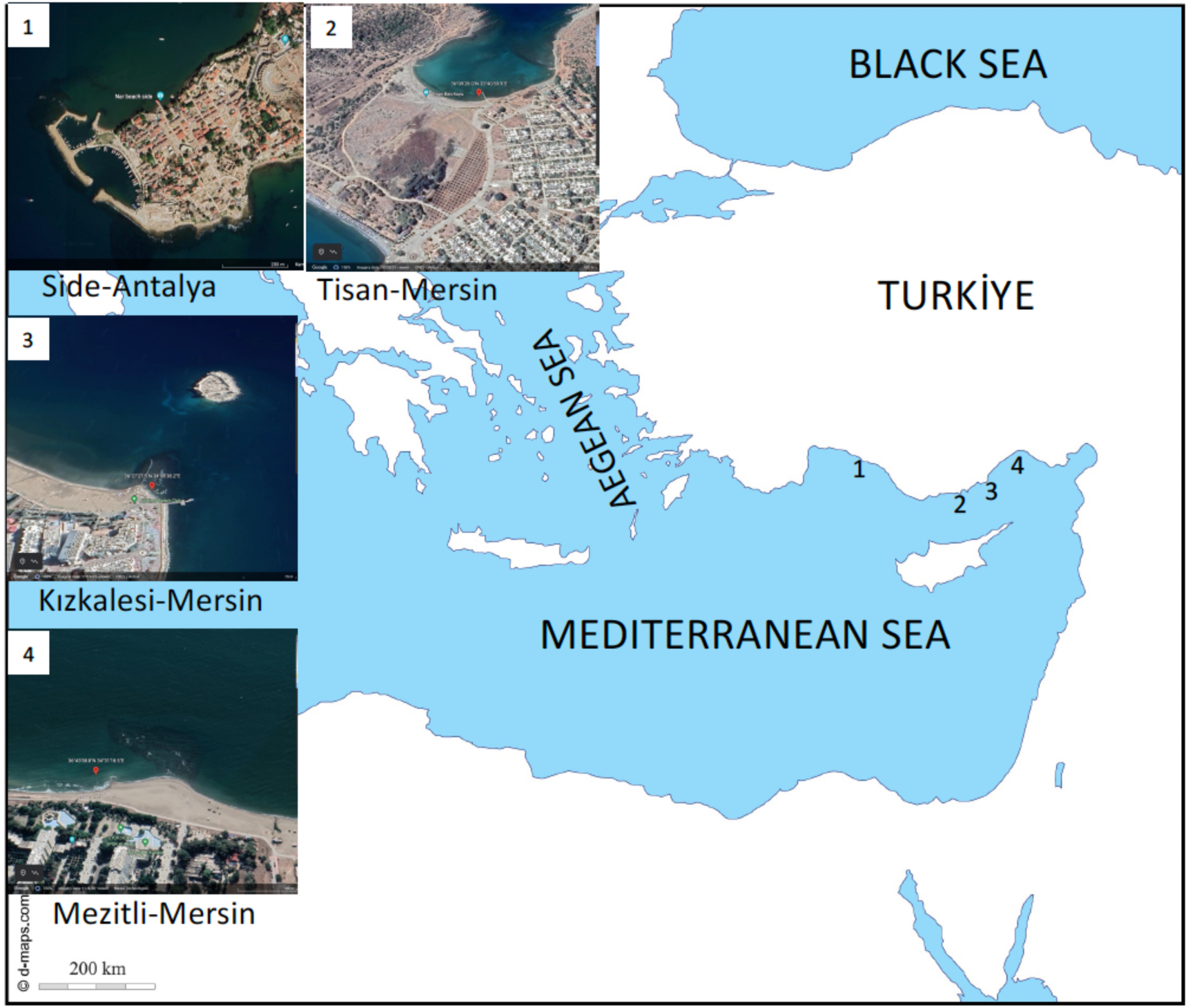
The sampling sites of the northeastern Mediterranean coastlines. Distance between the locations: Mezitli-Kızkalesi 72 km; Kızkalesi-Tisan 70 km; Tisan-Side 285 km.

The COI region was amplified using the Folmer *et al. (*1994) primer set (HCO2198; 5′-TAAACTTCAGGGTGACCAAAAAATCA-3′ and LCO1490; 5′- GGTCAACAAATCATAAAGATATTGG-3′), and the ITS2 region was amplified using White *et al*. (1990) ITS3-ITS4 primer set (5′-GCATCGATGAAGAACGCAGC-3′, 5′- TCCTCCGCTTATTGATATGC-3′). The PCR products were sequenced by Macrogen Inc. (Seoul, South Korea) in both directions.

In total, 81 COI and 11 ITS records of the order Suberitida were retrieved from GenBank, and SDA was performed alongside 11 samples from the present study. All sequences were aligned using MAFFT v7 (Katoh *et al*., 2017). The evolutionary distances between the samples from the present study and the database-mined samples were computed for both COI and ITS2 markers using the Kimura 2-Parameter model with MEGAX (v. 10.1) software. The correlation between geographic and genetic distances was tested using the Mantel Test (Mantel, 1967) on the BOLD system, comparing the geographic distance matrix (in kilometers) with the genetic divergence matrix.

### 2.1. Species delimitation analysis

After alignment and trimming, approximately 570 bp-length sequences were used for the analysis. We conducted SDA using two distinct approaches: the Automatic Barcode Gap Discovery method (also known as Assemble Species by Automatic Partitioning—ASAP) developed by Puillandre *et al*. (2021), which employs sequence similarity clustering, and the Poisson Tree Processes (PTP) method introduced by Zhang *et al*. (2013), which is based on tree-based coalescent processes. ASAP analyses were carried out using the web-based interface available at https://bioinfo.mnhn.fr/abi/public/asap/asapweb.html. We employed two metric options provided by ASAP for pairwise distance calculations: Jukes-Cantor (JC69) (Jukes and Cantor, 1969) and Kimura 2 parameter (K80) (Kimura, 1980). This method was selected to alleviate any possible bias arising from the choice of an evolutionary model for the delineation of Operational Taxonomic Units (OTUs).

PTP analyses (Zhang *et al.,* 2013) were conducted using the Bayesian implementation (bPTP), accessible through the web-based interface (http://species.hits.org/ptp/). For each alignment, a Bayesian tree was employed as input (Tang, 2014). These Bayesian trees were generated using the best-fit substitution models determined by Smart Model Selection (SMS) in PhyML (http://www.atgc-montpellier.fr/sms/execution.php). According to SMS, the optimal model for ITS2 was HKY85+G+I, while for COI, it was GTR+G+I. MrBayes v3.2 was utilized with these models. Two parallel analyses were conducted: 50,000 generations for ITS2 and 1,000,000 generations for COI. Trees were sampled every 100 generations, and the results of the initial 12,500 generations for ITS2 and 250,000 generations for COI were excluded as a burn-in fraction of 25% after confirming the stationarity of the lnL. Figtree (v1.4.4) was used to visualize the Bayesian tree.

## 3. Results

### 3.1. Morphological records

Samples exhibited radiant, smooth, jelly-like surfaces (gelatinoid), thin-encrusting textures, and dark blue colors (Figure 2A, C), which are characteristic features of *Terpios gelatinosus* (Bowerbank, 1866). Tylostyle types of macroscleres were observed in all samples (Figure 2B, D). The size range varied between approximately 94-330 µm for the samples from Kızkalesi, Mezitli, and Side (based on a total of 139 counted spicules, Figure 2B) and approximately 152-350 µm for the Tisan samples (based on a total of 61 counted spicules, Figure 2D).

**Figure 2.**
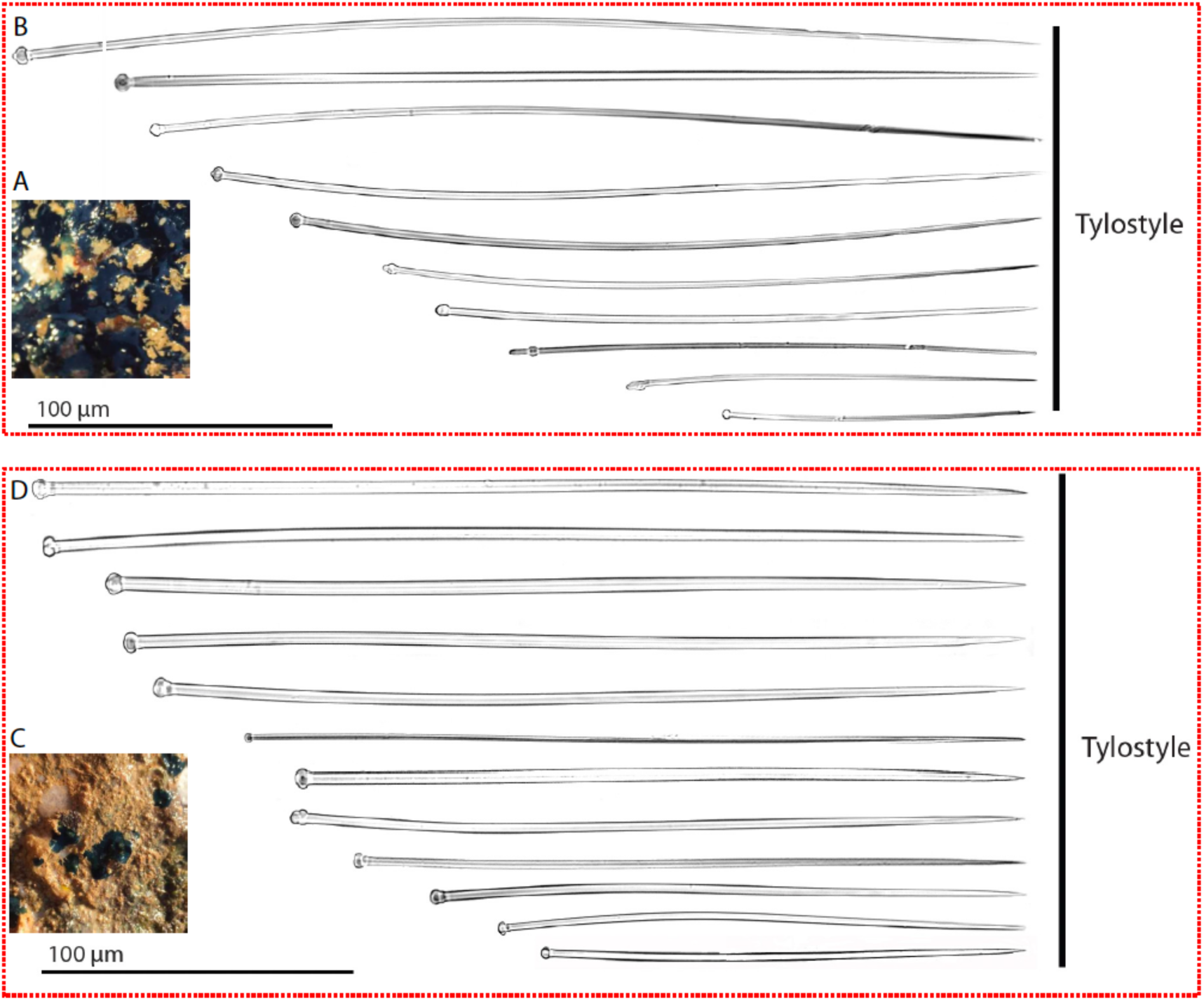
Basic morphologic records of *Terpios gelatinosus* from Kızkalesi **(A-B)** and. Tisan (**C-D**) regions. **A-C.** represent general images from the field and **B-D**. tylostyles under the light microscope (arranged according to sizes). Scale bars: 100 µm

### 3.2. Data deposition and Blast analysis results

All the sequences were uploaded to the Barcode of Life Data System (BOLD) and subsequently directed to NCBI, along with information on sampling areas, coordinates, and specimen images. Two different BOLD BIN IDs were assigned to the samples: ADY2734 for Tisan and AAK9893 for the samples from Side, Mezitli, and Kızkalesi. The NCBI accession numbers are OQ413306, OQ413307, OQ413308, OQ413309, OQ413310, OQ413311, OQ413312, OQ413313, and OQ413314.

Based on the BLAST results of the COI gene, the Kızkalesi, Mezitli, and Side samples exhibited 99-100% similarity to the *T. gelatinosus* species from Israel (KX866738.1, Idan et al., 2018) and *Protosuberites sp.* (AY561979.1, Nichols, 2005) from California, USA. Meanwhile, four Tisan samples showed a coverage of 97.4-97.6% (E-value 0.0) with the same species. The COI genetic distance between the two congeneric species, *Terpios hoshinota* and *T. gelatinosus*, ranged from 9.2% to 10.9% (Table 1). Unfortunately, due to the absence of an ITS2 reference sequence for *T. gelatinosus* in the database, no close matches were obtained in the BLAST analysis (Table 2). For the ITS2 region, the distance between the Tisan and Kızkalesi-Mezitli-Side samples was approximately 13%.

**Table 1.** Kimura two-parameter pairwise genetic distances (%) for the COI region of the *Terpios* genus. The samples with NCBI-ID were retrieved from GenBank. Present study sample’s location codes: K: Kızkalesi, M: Mezitli, S: Side, T: Tisan

| Sample name and location | 1 | 2 | 3 | 4 | 5 | 6 | 7 | 8 | 9 | 10 | 11 | 12 | 13 | 14 | 15 | 16 |
| --- | --- | --- | --- | --- | --- | --- | --- | --- | --- | --- | --- | --- | --- | --- | --- | --- |
| 1) *T. gelatinosus*_K |  |  |  |  |  |  |  |  |  |  |  |  |  |  |  |  |
| 2) *T. gelatinosus*_K | 0,2 |  |  |  |  |  |  |  |  |  |  |  |  |  |  |  |
| 3) *T. gelatinosus*_M | 0 | 0,2 |  |  |  |  |  |  |  |  |  |  |  |  |  |  |
| 4) *T. gelatinosus*_M | 0 | 0,2 | 0 |  |  |  |  |  |  |  |  |  |  |  |  |  |
| 5) *T. gelatinosus*_S | 0 | 0,2 | 0 | 0 |  |  |  |  |  |  |  |  |  |  |  |  |
| 6) *T. gelatinosus*_S | 0 | 0,2 | 0 | 0 | 0 |  |  |  |  |  |  |  |  |  |  |  |
| 7) *T. gelatinosus*_S | 0 | 0,2 | 0 | 0 | 0 | 0 |  |  |  |  |  |  |  |  |  |  |
| 8) *T. gelatinosus*_T | 2,4 | 2,6 | 2,4 | 2,4 | 2,4 | 2,4 | 2,4 |  |  |  |  |  |  |  |  |  |
| 9) *T. gelatinosus*_T | 2,4 | 2,6 | 2,4 | 2,4 | 2,4 | 2,4 | 2,4 | 0 |  |  |  |  |  |  |  |  |
| 10) *T. gelatinosus*_T | 2,4 | 2,6 | 2,4 | 2,4 | 2,4 | 2,4 | 2,4 | 0 | 0 |  |  |  |  |  |  |  |
| 11) *T. gelatinosus*_T | 2,4 | 2,6 | 2,4 | 2,4 | 2,4 | 2,4 | 2,4 | 0 | 0 | 0 |  |  |  |  |  |  |
| 12) KP764915.1-*T. hoshinota*-Indonesia | 9,3 | 9,2 | 9,2 | 9,2 | 9,2 | 9,2 | 9,2 | 10,7 | 10,7 | 10,7 | 10,7 |  |  |  |  |  |
| 13) MN507873.1 *T. hoshinota*-WA | 9,3 | 9,2 | 9,2 | 9,2 | 9,2 | 9,2 | 9,2 | 10,7 | 10,7 | 10,7 | 10,7 | 0,2 |  |  |  |  |
| 14) MN507872.1 *T. hoshinota*-WA | 9,5 | 9,5 | 9,5 | 9,5 | 9,5 | 9,5 | 9,5 | 10,9 | 10,9 | 10,9 | 10,9 | 0,4 | 0,4 |  |  |  |
| 15) GBMAA460-14 *T. hoshinota*-China | 9,5 | 9,5 | 9,5 | 9,5 | 9,5 | 9,5 | 9,5 | 10,9 | 10,9 | 10,9 | 10,9 | 0,4 | 0,4 | 0,6 |  |  |
| 16) KX866738.1 *T. gelatinosus*-Israel | 0 | 0,02 | 0 | 0 | 0 | 0 | 0 | 2,4 | 2,4 | 2,4 | 2,4 | 9,2 | 9,2 | 9,5 | 9,5 |  |
| 17) AY561979.1 *Protosuberites* sp. | 0,5 | 0,7 | 0,5 | 0,5 | 0,5 | 0,5 | 0,5 | 2,2 | 2,2 | 2,2 | 2,2 | 9,5 | 9,7 | 9,7 | 9,7 | 0,5 |

**Table 2.** Kimura two-parameter pairwise genetic distances (%) for the ITS region of the present study samples and *Suberites ficus.* Present study sample’s location codes: K: Kızkalesi, M: Mezitli, S: Side, T: Tisan

| Sample name and location | 1 | 2 | 3 | 4 | 5 | 6 | 7 | 8 | 9 | 10 |
| --- | --- | --- | --- | --- | --- | --- | --- | --- | --- | --- |
| 1) AJ627184.1 *Suberites ficus* |  |  |  |  |  |  |  |  |  |  |
| 2) *T. gelatinosus*_K | 17,5 |  |  |  |  |  |  |  |  |  |
| 3) *T. gelatinosus*_K | 17,5 | 0 |  |  |  |  |  |  |  |  |
| 4) *T. gelatinosus*_M | 17,5 | 0 | 0 |  |  |  |  |  |  |  |
| 5) *T. gelatinosus*_M | 17,5 | 0 | 0 | 0 |  |  |  |  |  |  |
| 6) *T. gelatinosus*_S | 17,5 | 0 | 0 | 0 | 0 |  |  |  |  |  |
| 7) *T. gelatinosus*_S | 17,5 | 0 | 0 | 0 | 0 | 0 |  |  |  |  |
| 8) *T. gelatinosus*_T | 21,2 | 12,8 | 12,8 | 12,8 | 12,8 | 12,8 | 12,8 |  |  |  |
| 9) *T. gelatinosus*_T | 21,2 | 12,8 | 12,8 | 12,8 | 12,8 | 12,8 | 12,8 | 0 |  |  |
| 10) *T. gelatinosus*_T | 21,2 | 12,8 | 12,8 | 12,8 | 12,8 | 12,8 | 12,8 | 0 | 0 |  |
| 11) *T. gelatinosus*_T | 21,2 | 12,8 | 12,8 | 12,8 | 12,8 | 12,8 | 12,8 | 0 | 0 | 0 |

### 3.3. Species delimitation for COI

In the SDA, the best score was determined using the ASAP method, where lower scores indicate better partitions (Puillandre *et al.,* 2021). The scores for the Kimura 2-parameter and Jukes-Cantor models were 3.5 and 4, respectively, resulting in the assignment of 23 OTUs (Figure 3). Additionally, genetic distances within the range of 0.005 to 0.05 were considered during the partitioning process. On the other hand, the PTP partition analysis assigned a total of 33 OTUs (Figure 4). The Tisan samples were assigned to different OTUs compared to the Kızkalesi-Mezitli-Side samples in the PTP analysis. In the ASAP analysis, all samples from the present study were grouped within the same OTU, along with 18 database samples representing *T. gelatinosus, Protosuberites sp., Protosuberites mereui, Halichondria sp., Pseudosuberites nudus,* and *Protosuberites denhartogi* (Figure 3). It’s worth noting that the congeneric species *T. hoshinota* was assigned to a different OTU by both the ASAP and PTP analyses, exhibiting approximately a 10% genetic distance from the samples in the present study (Figure 3, Table 1). The mean genetic distance among the 29 samples (Figure 3) was calculated to be 0.04, with pairwise distances ranging from 0.0 to 0.083.

**Figure 3.**
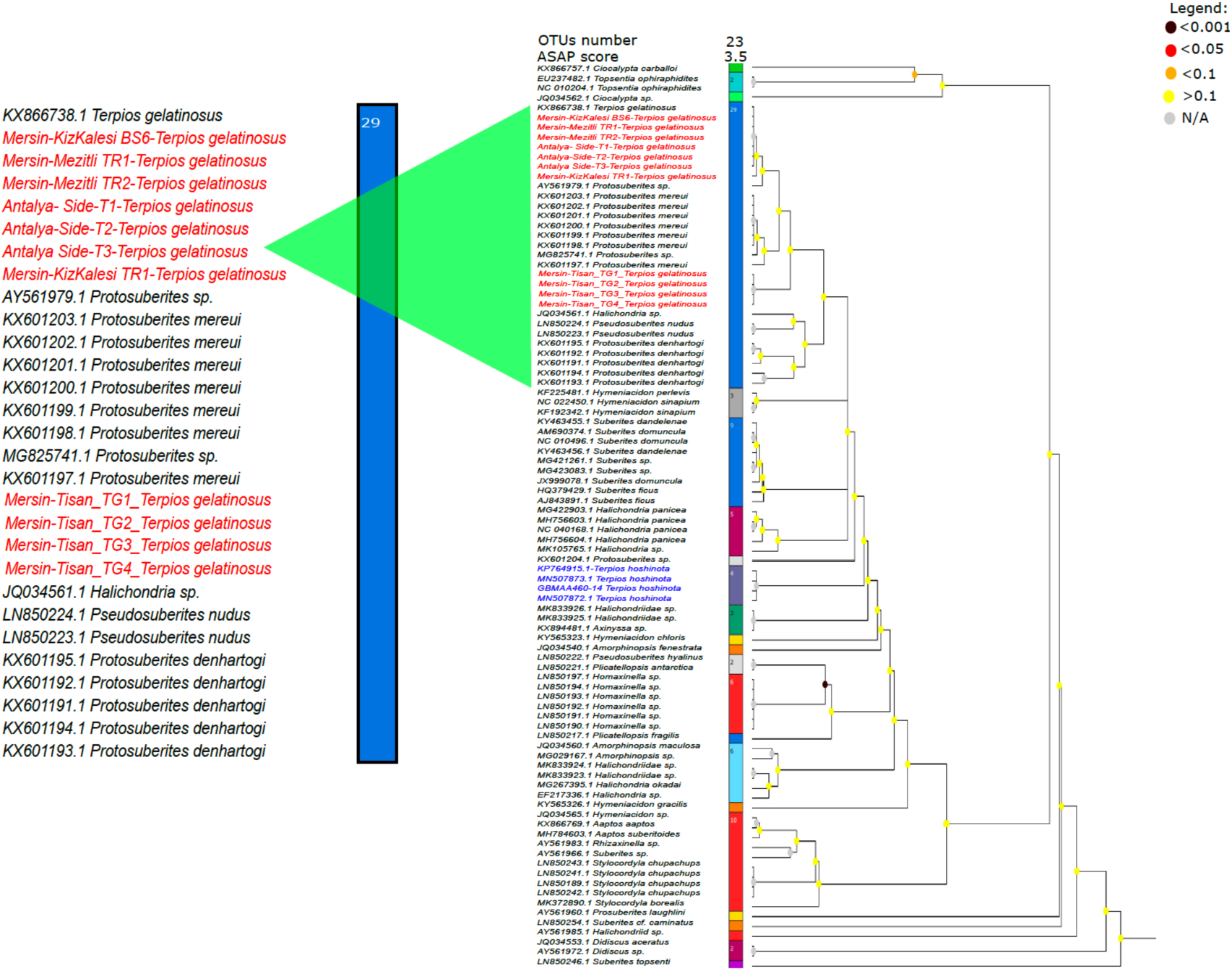
The figure displays the ASAP scores of Suberitida specimens’ COI. Different OTUs are represented by colors on the bar, and the number inside each color bar corresponds to the assigned specimen number for that OTU. The number of subsets, assigned total OTU number, ASAP score (where lower scores indicate better partitions, Puillandre *et al.,* 2021), and the best rank column (1) are included. Samples from the present study are marked in red letters. Legend: Darker colors in the figure indicate lower probabilities, while a gray dot signifies that the probability was not computed. When a probability is very low (dark color), it suggests that the groups within the node likely correspond to different species. The blue letters represent the congeneric species, *Terpios hoshinota*.

**Figure 4.**
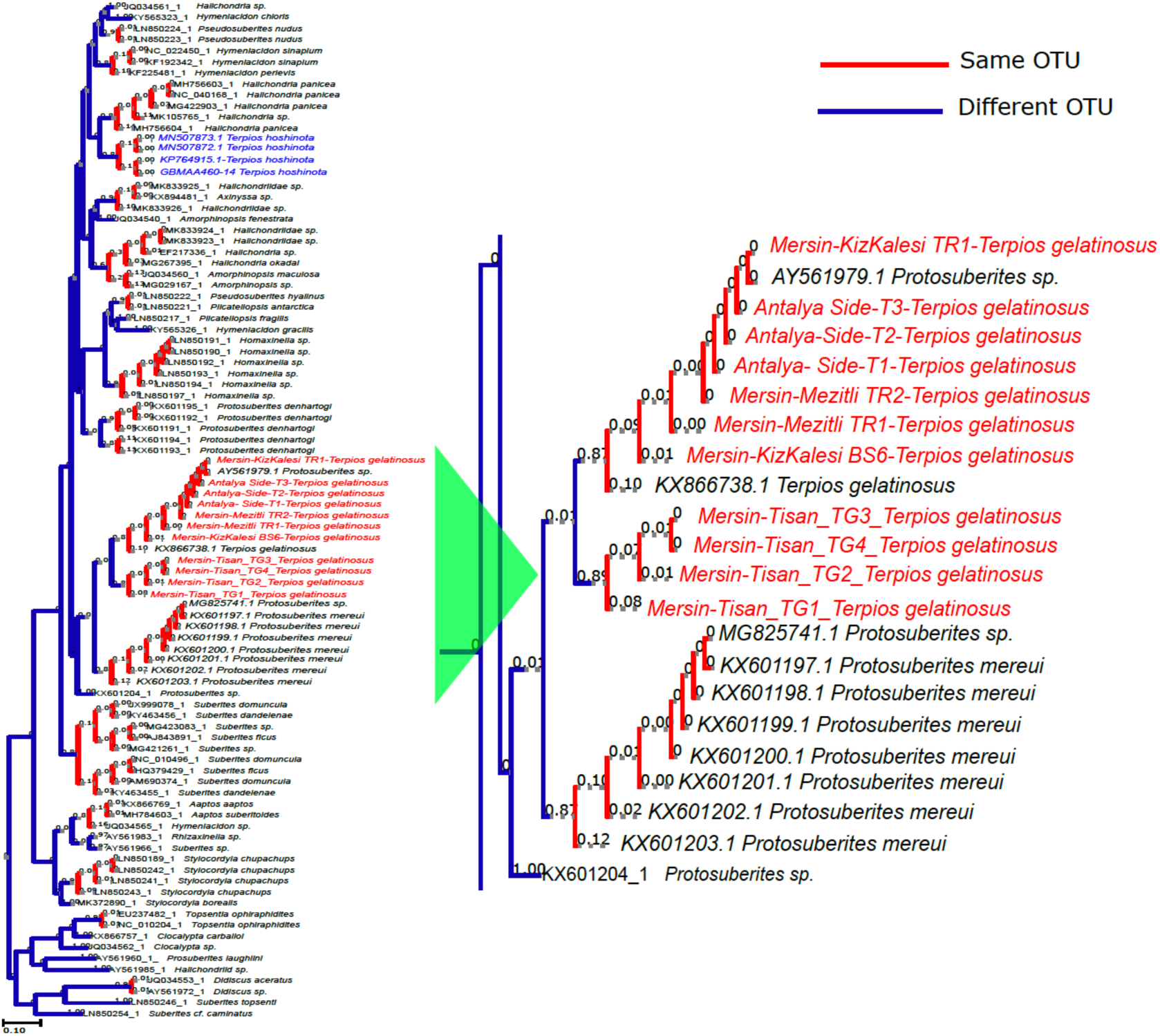
PTP score of Suberitida specimens’ COI: blue lines represent different OTUs, red lines represent the same OTUs. The red letter represents present study samples. The blue letters represent the congeneric species, *Terpios hoshinota*.

### 3.4. Mantel Test

According to the Mantel test, no correlation was recorded between the geographic distance and the genetic divergence (Rsq=0.001, P-value =0.37).

### 1.1. Species delimitation for ITS2

The aligned and trimmed sequence length was 240 bp. In the SDA, the lowest ASAP score was 1.5 for both the Kimura 2-parameter and Jukes-Cantor models. As a result, eleven OTUs were assigned by both the ASAP (Figure 5A) and PTP (Figure 5B) analyses. In these analyses the Kızkalesi-Mezitli-Side samples were grouped within one OTU, while the Tisan samples were in another OTU. The genetic distance between these two OTUs was approximately 13% (Table 2).

**Figure 5.**
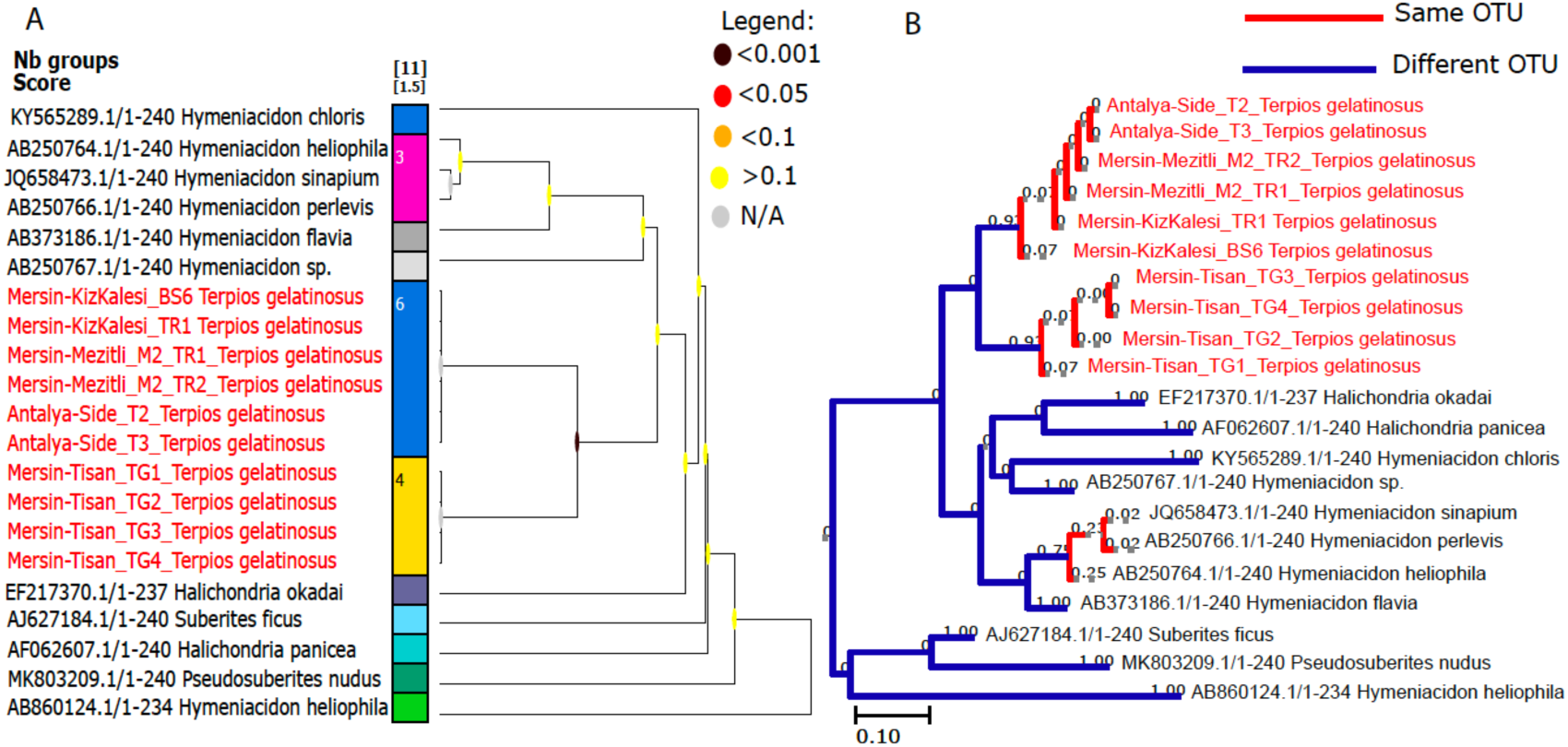
**A)** The figure displays the ASAP scores of Suberitida specimens’ ITS. Different OTUs are represented by colors on the bar, and the number inside each color bar corresponds to the assigned specimen number for that OTU. The number of subsets, assigned total OTU number, ASAP score (where lower scores indicate better partitions, Puillandre *et al.,* 2021), and the best rank column (1) are included. Samples from the present study are marked in red letters. Legend: Darker colors in the figure indicate lower probabilities, while a gray dot signifies that the probability was not computed. When a probability is very low (dark color), it suggests that the groups within the node likely correspond to different species. **B)** PTP score of Suberitida specimens’ ITS; blue lines represent different OTUs, red lines represent the same OTUs. The red letter represents present study samples.

### 3.6. Species Assignment

Species assignment was based on a combination of basic morphological data, BLAST scores, ASAP, and PTP analyses. Despite minor variations in spicule sizes, both morphological analysis and COI ASAP scores suggested the presence of a single species (*T. gelatinosus*) in the area. However, the COI PTP scores indicated the presence of two distinct species. Furthermore, ITS results from both ASAP and PTP analyses suggested the potential existence of cryptic speciation in the area.

## 2. Discussion

Recording species distribution and identifying new species are crucial aspects of scientific research. These endeavors significantly contribute to our understanding of ecology, evolution, inter-organism relationships, biodiversity conservation, and even potential biomedical applications (Mora *et al*., 2011). In this study, we unveiled the presence and spatial distribution of the *T. gelatinosus* species, along with indications of possible cryptic speciation, in the shallow waters along the northeastern Mediterranean coastlines. Our approach involved utilizing basic morphological data, a distance matrix incorporating two molecular markers, and scores from PTP and ASAP analyses for species identification.

Both COI and ITS markers are frequently used for species delimitation. Each marker offers distinct advantages and disadvantages depending on the research objectives and the organisms under investigation. COI, with its rapid evolutionary rate, is conserved across a broad spectrum of taxa, making it a valuable tool for identifying species in diverse groups (Hebert *et al*., 2003b). Using the ITS region, on the other hand, is advantageous due to its high variability, which enables the differentiation of closely related species when used alongside other data, such as morphological characteristics or additional genetic markers (Wörheide *et al*., 2002; Erpenbeck *et al*., 2006, 2007; Erwin & Thacker *et al.,* 2007). While a 2.5% divergence for the COI marker is typically accepted as a threshold for most eukaryotes, it is important to note that different taxa, including Porifera, can exhibit varying divergence ranges due to their unique evolutionary histories (Hajibabaei *et al*., 2007; Núñez-Pons *et al*., 2017). Therefore, the ITS region has been proposed as a potential alternative to the COI barcode region for sponges, attributed to their low evolutionary pressure and high variation rates (Wörheide & Erpenbeck, 2007; Song *et al*., 2012). In the present study, while the morphological data and the ASAP score for COI indicated no discrepancies, the PTP score for COI, along with both ASAP and PTP scores for the ITS marker, suggested the potential presence of cryptic speciation in the studied areas.

Cryptic speciation, involving the existence of two or more species that closely resemble each other but are genetically distinct and unable to interbreed, can arise due to various factors such as geographical isolation, habitat differentiation, or ecological adaptation differences (Nosil & Crispi, 2006; Bickford *et al.,* 2007; Worsham *et al.,* 2017). Identifying and distinguishing cryptic species can be challenging due to their similar external appearances. However, genetic techniques are increasingly being employed to uncover hidden biodiversity in sponges and other organisms (Hooper & van Soest, 2002). Despite low genetic divergence, cryptic speciation has been reported for the *Hemimycale columella* species through the utilization of genetic markers such as COI, 18S, and 28S gene regions (Uriz *et al*., 2017). On the other hand, COI and ITS markers were used to elucidate the morphospecies complex of *Lanthella basta* and identify cryptic speciation in northern Australia (Andreakis *et al*., 2012). In our case, we used COI and ITS to reveal potential cryptic speciation in the Tisan region.

Many cryptic sponge species, which are geographically separated, are believed to have undergone allopatric speciation (Uriz & Turon, 2012). Sponges are sessile organisms with limited dispersal capabilities during their juvenile stage. Consequently, gene flow between sponge populations is influenced not only by larval dispersal but also by geographical distance (Maldonado, 2006). In our study, we conducted a Mantel test and found no significant correlation between genetic connectivity and geographic distance. Interestingly, both COI’s PTP analysis and ITS’s ASAP-PTP analyses revealed that the Kızkalesi-Mezitli-Side samples were grouped under a different OTU compared to the Tisan samples, despite the Tisan site being located between the Kızkalesi and Side sites. This unexpected distribution pattern may be explained by marine traffic, which plays a role in the dispersal of this organism. The Kızkalesi-Mezitli-Side sites are situated near heavy shipping routes (Alexopoulos, 2013), while the Tisan site is a relatively protected area, distant from ports. This leads us to consider the possibility of allopatric cryptic speciation in the studied area.

On the other hand, *Terpios fugax* is one of the closest relatives of *T. gelatinosus*. It has been recorded in the Atlantic Ocean (de Laubenfels and Hindle, 1950), the Indian Ocean (Pattanayak, 1999), the Pacific Ocean (de Laubenfels, 1954), and the European Seas (Voultsiadou-Koukoura & van Soest, 1993). According to Araya and Rützler (2016), all European records of *T. fugax* are most likely misidentified individuals of *T. gelatinosus*. However, we could not compare our samples to *T. fugax* because there is no public DNA record of this species in any database.

This study constitutes the documentation of *T. gelatinosus* throughout the 450 km eastern Mediterranean region. The findings from the SDA, which utilized two DNA markers alongside basic morphological data, strongly indicate the potential for cryptic speciation along the Mediterranean coasts of Türkiye. Following a comprehensive morphological examination, it is possible that the Tisan samples may be classified as a different species within the *Terpios* genus. Additionally, the enhanced resolution within the *Terpios* genus makes the ITS region a strong candidate for a potential barcode site for Suberitidae species. In addition to this primary finding, our research has demonstrated that the two closely related species, *T. gelatinosus* and *T. hoshinota*, are located on separate branches; similar outcomes for other sponge species were also documented (Gazave *et al.,* 2010; Morrow *et al.,* 2012), this could indicate a potential polyphyletic origin for the *Terpios* genus.

## Acknowledgments

We would like to express our sincere thanks to Erdemli and Manavgat municipalities, as well as to Fatima Nur Oğul and Begüm Ece Tohumcu for their invaluable assistance during the sampling process.

## Authors’ contributions

A.K. conceived and planned research; E.Ö., B.T., and A.K., carried out the experiments; E.Ö., B.T., and A.K. wrote the manuscript.

## Financial support

This study was financially supported by Middle East Technical University (grant number YÖP-701-2018-2666) and the IMS-METU DEKOSIM project (grant number BAP-08-11-DPT2012K120880).

## Availability of data and material

The datasets generated during and/or analyzed during the current study are available in the BOLD system BOLD (BIN IDs: ADY2734 and AAK9893) and NCBI (for COI: OQ413306, OQ413307, OQ413308, OQ413309, OQ413310, OQ413311, OQ413312, OQ413313, and OQ413314, for ITS OQ507665, OQ507666, OQ507667, OQ507668, OQ507669, OQ507670, OQ507671, OQ507672, OQ507673, OQ507674, OQ507675)

## Competing of interest

The authors declare that they have no conflict of interest.

## Ethics approval

All applicable international, national, and/or institutional guidelines for the care and use of animals were followed.

## Notes

### Competing Interest Statement

The authors have declared no competing interest.

### Summary of Updates

References in both the in-text citations and reference section have been revised. Five references were replaced. Grammatical errors have been corrected, and the table formatting has been updated.

